# Stripes and loss of color in ball pythons (*Python regius*) are associated with variants affecting endothelin signaling

**DOI:** 10.1101/2022.10.09.511500

**Authors:** Uyen M. Dao, Izabella Lederer, Ray L. Tabor, Basmah Shahid, The BIO306W Consortium, Chiron W. Graves, Hannah S. Seidel

## Abstract

Color patterns in non-avian reptiles are beautifully diverse, but little is known about the genetics and development of these patterns. Here we investigated color patterning in pet ball pythons (*Python regius*), which have been bred to show color phenotypes that differ dramatically from the wildtype form. We report that several color phenotypes in pet animals are associated with putative loss-of-function variants in the gene encoding endothelin receptor EDNRB1: (i) frameshift variants in *EDNRB1* are associated with conversion of the normal mottled color pattern to skin that is almost fully white, (ii) missense variants affecting conserved sites of the EDNRB1 protein are associated with dorsal, longitudinal stripes, and (iii) substitutions at *EDNRB1* splice donors are associated with subtle changes in patterning compared to wildtype. We propose that these phenotypes are caused by loss of specialized color cells (chromatophores), with loss ranging from severe (fully white) to moderate (dorsal striping) to mild (subtle changes in patterning). Our study is the first to describe variants affecting endothelin signaling in a non-avian reptile and suggests that reductions in endothelin signaling in ball pythons can produce a variety of color phenotypes, depending on the degree of color cell loss.

## Introduction

Skin colors in vertebrates originate from specialized color cells. These cells are known as chromatophores, and they come in three main types (Mills and Patterson 2009; Kelsh *et al*. 2009; Cuthill *et al*. 2017). Melanophores synthesize the pigment melanin, whose color ranges from black to reddish brown. Xanthophores store red-to-yellow pigments, including pteridines synthesized by the cell and carotenoids obtained from the diet. Iridophores contain platelets of crystallized purines, which can appear blue, white, or iridescent, depending on their structure. Skin color in non-avian reptiles and lower vertebrates is determined by layering of different chromatophore types (Patterson and Parichy 2019; Kuriyama *et al*. 2020). Green colors, for example, are produced by yellow xanthophores above blue iridophores (e.g. Saenko *et al*. 2013). Golden brown colors, as a second example, are produced by a combination of brown melanophores, yellow xanthophores, and white iridophores (e.g. Brejcha *et al*. 2019). Mammals possess only one type of chromatophore (melanophores), and their skin color is limited to shades of brown and black (Mort *et al*. 2015).

Chromatophores are derived from the neural crest and migrate during development to reach their final positions in the skin (Parichy and Spiewak 2015; Mort *et al*. 2015; Haupaix and Manceau 2020). Once in the skin, chromatophores organize into patterned arrangements to produce the overall color pattern of the animal. This process has been best characterized in zebrafish, where patterning is driven by short- and long-range interactions among chromatophores themselves (Singh and Nuesslein-Volhard 2015; Patterson and Parichy 2019). Similar mechanisms likely function in mammals to produce periodic markings such as stripes and spots (Kaelin *et al*. 2012, 2021; Mallarino *et al*. 2016). Pattern formation is less well understood in non-avian reptiles, although the idea that similar mechanisms may function in this group is supported by theoretical studies showing that Turing-like interactions can produce many of the basic color patterns observed in this group (Murray and Myerscough 1991; Kondo and Shirota 2009; Kuriyama *et al*. 2020; Ascarrunz and Sanchez-Villagra 2022).

An emerging model for understanding coloration and pattern formation in non-avian reptiles is the ball python (*Python regius*) (Garcia-Elfring *et al*. 2020; Brown *et al*. 2022). Ball pythons are native to west Africa, but have become common as pets in the United States. Wild ball pythons exhibit a mottled color pattern, consisting of irregular black and golden-brown patches of skin (Fig 1). Pet ball pythons, by contrast, have been bred to include animals with dramatically different color patterns (McCurley 2005; Broghammer 2019). These variants, known as ‘color morphs,’ include animals with solid body colors, longitudinal stripes, or major changes in pattern patchiness. Many of these phenotypes are reminiscent of patterning changes that have occurred within the snake lineage across evolutionary time, including patterns thought to be adaptive for certain behavioral ecologies (Brodie 1992; Olsson *et al*. 2013; Allen *et al*. 2013; Pizzigalli *et al*. 2020). Color morphs in ball pythons may therefore provide insight into the genetic and developmental mechanisms controlling patterning changes across species.

**Fig 1.**
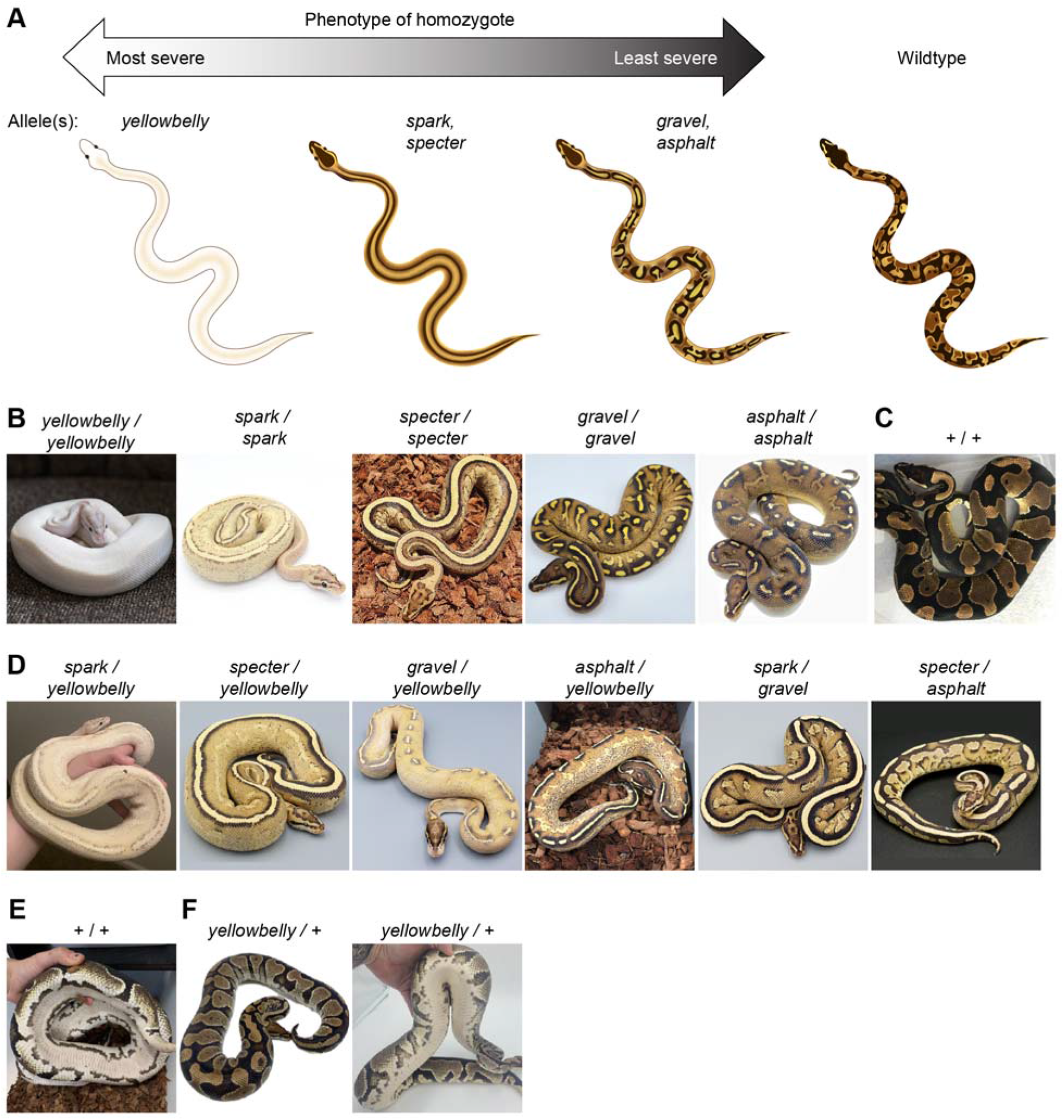
Phenotypes of color morphs in the Yellowbelly series. (A) Schematic of homozygote phenotypes in the Yellowbelly series. (B) Ball pythons described as homozygous for each allele in the Yellowbelly series. (C) Wildtype ball python. (D) Ball pythons described as compound heterozygous for alleles in the Yellowbelly series. (E) Underbelly of a wildtype ball python. (F) Ball python described as heterozygous for the *yellowbelly* allele. The underbelly is subtly lighter and differently patterned compared to wildtype. (B-F) Images are representative examples. Variation exists within each genotype. Photos, Chiron Graves, Gavin Costello, Ball Python Shed, Ball In Hand Pythons, Barbara Penyak, Freedom Breeder, Patrick Faulkner, Off The Wall Balls, Colorado Cold Bloods, Laura Carter, Brent McKelvey, Travis Wyman.

Many color morphs in ball pythons show simple dominant or recessive patterns of inheritance. These inheritance patterns suggest that these color morphs are caused by variants in single genes (e.g. Garcia-Elfring *et al*. 2020; Brown *et al*. 2022). Some morphs belong to allelic series, with some morphs in the series showing more dramatic phenotypes than others (McCurley 2005; Broghammer 2019). Such morphs are thought to represent different molecular variants of the same gene, akin to mutants in allelic series arising from classical mutagenesis screens in model organisms.

One example of an allelic series in ball pythons is the Yellowbelly series of color morphs (Fig 1). This series is described by breeders as having five alleles (*yellowbelly, spark, specter, gravel*, and *asphalt*). Morphs in the Yellowbelly series exhibit changes in patterning and reduced coloration compared to wildtype. These phenotypes range from severe to mild (Fig 1A and 1B). Homozygotes of the strongest allele (*yellowbelly*) have skin that is almost fully white (Fig 1B). The intermediate alleles (*spark* and *specter*) produce homozygotes with a pair of dorsal, longitudinal stripes (Fig 1B). These animals also have moderately reduced coloration. The weakest alleles (*gravel* and *asphalt*) produce homozygotes with subtle alterations in patterning and mildly reduced coloration (Fig 1B). Compound heterozygotes show phenotypes intermediate between the parents, often including dorsal, longitudinal stripes (Fig 1D). These phenotypes are largely recessive, and heterozygotes of any of these alleles in combination with the wildtype allele show color patterns that are largely similar to wildtype (Fig 1F). Some heterozygotes have slightly lighter underbellies than wildtype (Figs 1E and 1F), but this phenotype is variable and difficult to recognize, especially for alleles other than the *yellowbelly* allele.

All-white skin in ball pythons is not the outcome of a defect in pigment synthesis. Such defects remove a single pigment (e.g. melanin) without affecting other pigments (e.g. Brown *et al*. 2022). Instead, all-white skin suggests a loss of multiple types of chromatophores. The absence of brown-to-black coloration suggests an absence of melanophores, and the absence of yellow coloration suggests an absence of xanthophores. Iridophores in animals with all-white skin are harder to asses, given that iridophores in ball pythons are likely white-reflecting and thus cannot be detected by visual appearance alone. In all-white morphs of other reptiles and amphibians, for example, iridophores are present in some cases (e.g. Ullate-Agote and Tzika 2021) and absent in others (e.g. Woodcock *et al*. 2017). Loss of chromatophores has been described in other vertebrates, where the underlying defects include lack of chromatophore specification, insufficient proliferation of chromatophore precursors, and failure of these precursors to migrate from the neural crest to their normal positions in the skin (reviewed in Mort *et al*. 2015; Patterson and Parichy 2019).

A prominent phenotype in the Yellowbelly series is conversion of the wildtype color pattern to dorsal, longitudinal stripes (Fig 1B). We hypothesized that this phenotype, like all-white skin, was the outcome of chromatophore loss, albeit to a lesser degree. This hypothesis arises from a resemblance between dorsal striping in ball pythons and a mutant phenotype in zebrafish caused by fewer chromatophores. The adult color pattern in zebrafish is initiated by iridophore precursors that migrate via the horizontal midline to populate the skin (Fig 2A) (Singh and Nuesslein-Volhard 2015; Patterson and Parichy 2019). In mutant zebrafish with fewer iridophores, iridophores fail to populate the skin and instead remain clustered at the horizontal midline (Parichy *et al*. 2000; Lopes *et al*. 2008; Patterson and Parichy 2013; Frohnhöfer *et al*. 2013; Mo *et al*. 2017; Spiewak *et al*. 2018). Melanophores in zebrafish require iridophores for survival, and hence melanophores in these mutants are less abundant and persist only near iridophores (Patterson and Parichy 2013; Frohnhöfer *et al*. 2013; Patterson *et al*. 2014). The resulting phenotype is a pair of midline stripes on each side of the fish’s body (Fig 2B). We suspected that these midline stripes might be analogous to dorsal stripes in the Yellowbelly series, given that chromatophores in ball pythons likely migrate from the neural crest in a dorsal-to-ventral path, as they do in other higher vertebrates (Mort *et al*. 2015). Incomplete loss of chromatophores in ball pythons might therefore cause chromatophores to remain clustered near the dorsal ridge. This hypothesis provides a simple explanation for the range of phenotypes observed in the Yellowbelly series: all-white skin may be outcome of chromatophore loss that is severe (Fig 2F); dorsal striping may be the outcome of chromatophore loss that is moderate (Fig 2E); and subtle alternations in patterning may be the outcome of chromatophore loss that is mild (Fig 2D). The cellular basis of this loss could plausibly include defects in chromatophore specification, proliferation, or migration, or a combination thereof.

**Fig 2.**
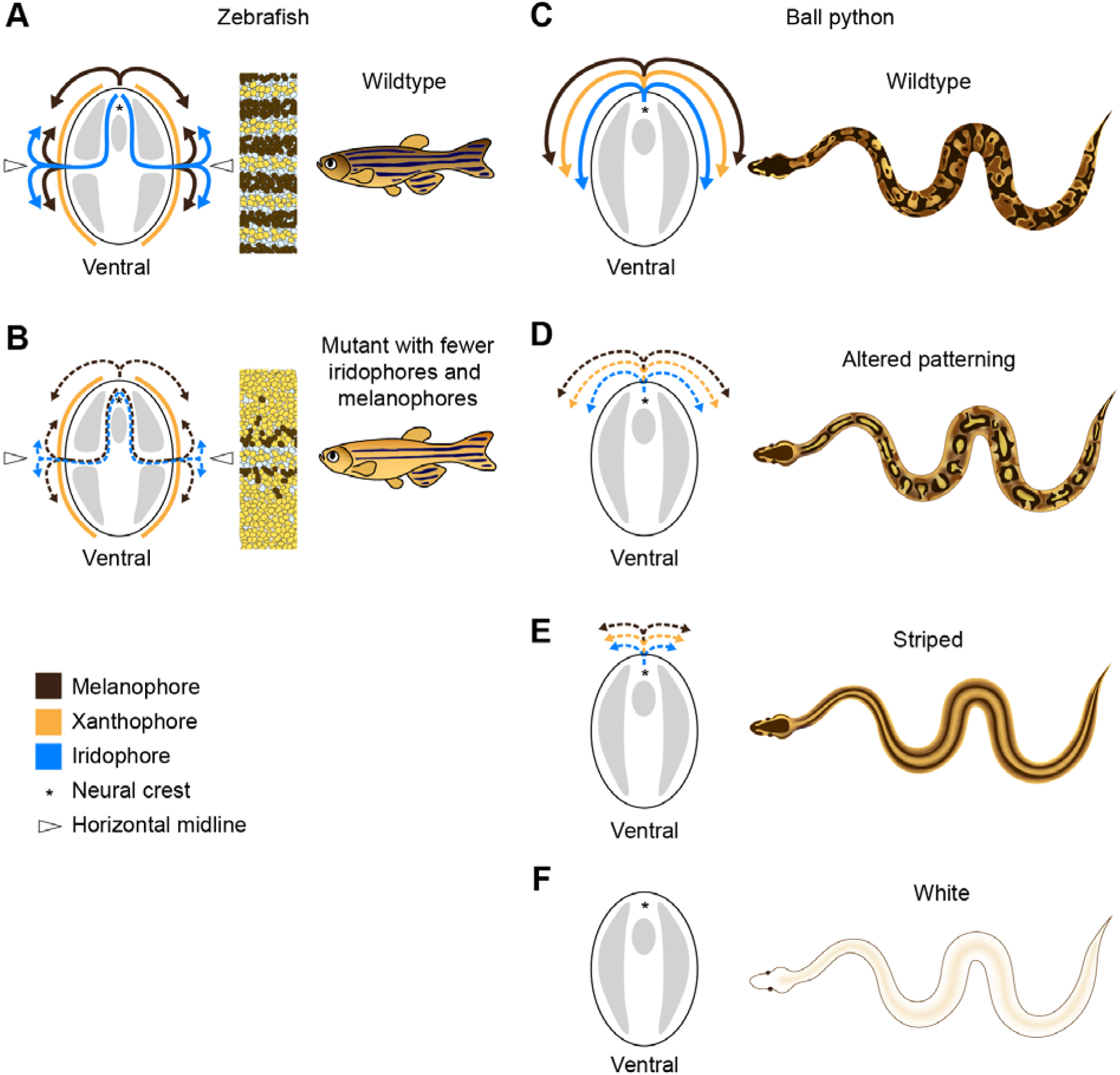
Phenotypes in the Yellowbelly series are hypothesized to arise from loss of chromatophores. (A) Simplified model of chromatophore migration and stripe formation during the larval-to-adult transition in zebrafish (Singh and Nuesslein-Volhard 2015; Patterson and Parichy 2019). Xanthophore precursors (yellow) persist in the epidermis from larvalhood. Melanophore and iridophore precursors (brown and blue) migrate from the neural crest to the epidermis along nerve tracks. The primary route for iridophores is through the horizontal midline. The adult pattern consists of dark stripes (melanophores and sparse iridophores) and light inter-stripes (xanthophores and dense iridophores). (B) Schematic of mutant zebrafish with a primary loss of iridophores and a secondary loss of melanophores (Parichy *et al*. 2000; Lopes *et al*. 2008; Patterson and Parichy 2013; Frohnhöfer *et al*. 2013; Mo *et al*. 2017; Spiewak *et al*. 2018). Iridophores fail to populate the skin and remain clustered near the horizontal midline. Melanophores require iridophores for survival and persist only near the iridophores. (C) Hypothesized model for chromatophore migration in ball pythons. Chromatophores are presumed to migrate in a dorsal-to-ventral path, as they do in other higher vertebrates (Mort *et al*. 2015). (D-F) Phenotypes in the Yellowbelly series are hypothesized to be caused by a loss of chromatophores. Loss is hypothesized to range from mild (altered patterning) to moderate (striped) to severe (white). This model depicts a loss of all three types of chromatophores, but the loss of iridophores is uncertain, and the range of developmental processes plausibly affected includes chromatophore specification, proliferation, migration, or a combination thereof.

Our hypothesis about chromatophore loss in the Yellowbelly series suggested candidate genes that might be causative for these phenotypes. At the top of this list were genes encoding components of endothelin signaling. Endothelin signaling comprises a vertebrate-specific signaling pathway responsible for regulating the development of neural crest cells (Saldana-Caboverde and Kos 2010; Bondurand *et al*. 2018). Components of endothelin signaling include endothelin ligands (endothelins), G-protein coupled receptors (endothelin receptors), and enzymes responsible for processing the ligands (endothelin-converting enzymes). Most vertebrate genomes encode three endothelin ligands (EDN1, EDN2, and EDN3) and three endothelin receptors (EDNRA, EDNRB1, and EDNRB2) (Braasch *et al*. 2009). Exceptions include zebrafish and eutherian mammals, which have lost *EDNRB2* (Braasch *et al*. 2009). EDNRA is selective for ligands EDN1 and EDN2 and helps pattern the skeleton of the face and head (Krystek *et al*. 1994; Lee *et al*. 1994; Lecoin *et al*. 1998; Liu *et al*. 2019; but see Menzi *et al*. 2016; Clouthier *et al*. 2010). EDNRB1 and EDNRB2, by contrast, regulate chromatophore precursors, among other cell types. EDNRB1 is required for proper proliferation and migration of chromatophore precursors in zebrafish and mammals, and EDNRB2 plays a similar role in birds (e.g. Baynash *et al*. 1994; Parichy *et al*. 2000; Pla *et al*. 2005; Harris *et al*. 2008; reviewed in Braasch and Schartl 2014). EDNRB1 and EDNRB2 are activated in cell culture by all three endothelin ligands (Krystek *et al*. 1994; Lecoin *et al*. 1998; Liu *et al*. 2019), but their primary ligand *in vivo* is thought to be EDN3. Loss of EDN3 phenocopies loss of EDNRB1 in zebrafish and mammals (Baynash *et al*. 1994; Edery *et al*. 1996; Hofstra *et al*. 1996; Bondurand *et al*. 2006; Pingault *et al*. 2010; Spiewak *et al*. 2018), and reduced *EDN3* expression causes similar phenotypes in amphibians (Kawasaki-Nishihara *et al*. 2011; Woodcock *et al*. 2017). EDN3 is a potent mitogen of chromatophore precursors across vertebrates (Lahav *et al*. 1996; Reid *et al*. 1996; Dupin *et al*. 2000), and increased *EDN3* expression is associated with darker skin and hair in birds and mammals (Dorshorst *et al*. 2011; Shinomiya *et al*. 2012; Kaelin *et al*. 2021). *EDNRB1, EDNRB2*, and *EDN3* therefore play conserved roles in chromatophore development in vertebrates and are candidate genes for the causative gene of the Yellowbelly series.

The goal of the current study was to identify the genetic cause of color phenotypes in the Yellowbelly series. We predicted that these phenotypes were caused by loss-of-function variants in the endothelin ligand gene *EDN3* or the endothelin receptor genes *EDNRB1* or *EDNRB2*. Our approach was to recruit samples of pet ball pythons (shed skins) from across the United States and search for putative loss-of-function variants in these genes.

## Results

### The *yellowbelly* allele is associated with frameshift variants in *EDNRB1*

We hypothesized that morphs in the Yellowbelly series were caused by loss-of-function variants in *EDN3, EDNRB1*, or *EDNRB2*. To test this hypothesis, we amplified and sequenced the coding regions and adjacent splice sites of each gene in one ball python described as a *yellowbelly* homozygote by its owner. Sequences from this animal were compared to sequences from an animal having normal coloration (henceforth ‘wildtype’). We observed that the *yellowbelly* homozygote was homozygous for a 1-bp deletion in the fourth coding region of *EDNRB1* (OP589186:c.1646del, Fig 3B). No coding or splice-site variants were observed in *EDN3* or *EDNRB2*. The 1-bp deletion in *EDNRB1* introduces a frameshift in the transcript, which removes or alters three of the seven transmembrane helices of the EDNRB1 protein (Fig 3E). This truncation likely abolishes protein function, given that G-protein-coupled receptors are generally unable to function without an intact transmembrane structure (Rosenbaum *et al*. 2009). We conclude that the 1-bp deletion is likely a strong loss-of-function allele of *EDNRB1*.

**Fig 3.**
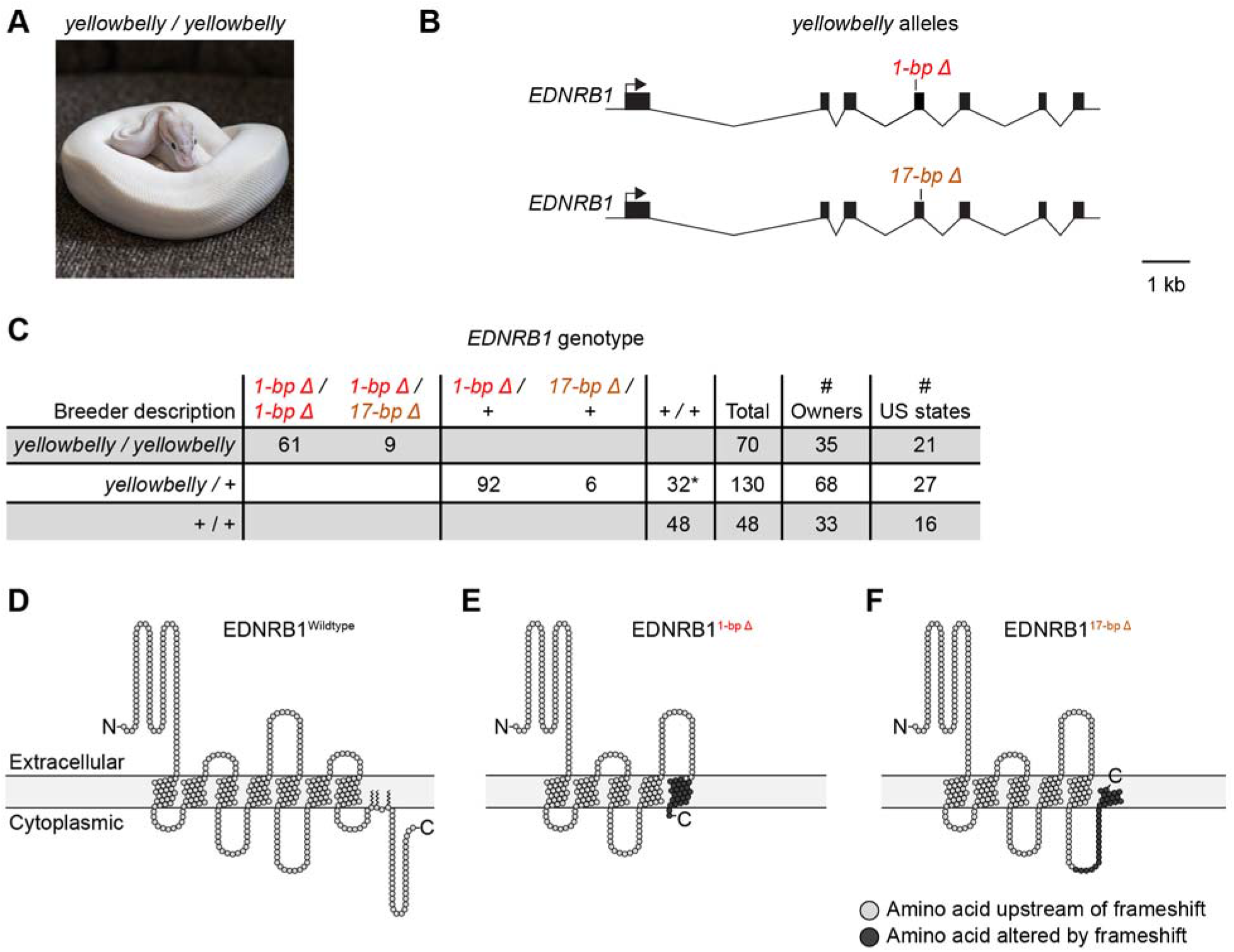
The *yellowbelly* allele is associated with frameshift variants in *EDNRB1*. (A) Ball python described as homozygous for the *yellowbelly* allele. Photo, Barbara Penyak. (B) Schematic of molecular *EDNRB1* alleles associated with the *yellowbelly* allele. One allele contains a 1-bp deletion. The other allele contains a 17-bp deletion. (C) Table showing association between the *yellowbelly* allele and the 1-bp and 17-bp deletions. This table includes the original *yellowbelly* homozygote in which the 1-bp deletion was identified. Numbers of owners and US states indicate the total numbers of owners and US states from which the animals were collected. *, presumed mis-labeling by owner, including one animal heterozygous for *L152F* (see Discussion). (D) Schematic of wildtype EDNRB1 protein, created using Protter (Omasits *et al*. 2014). (E-F) Schematics of truncated EDNRB1 proteins resulting from the 1-bp and 17-bp deletions. Schematics are intended to show premature terminations and do not imply that the truncated proteins fold properly and are inserted into the cell membrane.

To test whether the 1-bp deletion in *EDNRB1* was associated with phenotypes attributed to the *yellowbelly* allele, we genotyped 117 additional animals for this deletion. These animals consisted of 69 animals described as *yellowbelly* homozygotes and 48 animals described as having normal coloration or belonging to morphs other than those in the Yellowbelly series (henceforth ‘Non-Yellowbelly’ animals). These animals were collected from a total of 35 and 33 pet owners, respectively, and thus represent a broad sampling of the pet population. We observed that all animals described as *yellowbelly* homozygotes were either homozygous for the 1-bp deletion (60 animals) or heterozygous for it (nine animals) (Fig 3C). By contrast, none of the Non-Yellowbelly animals carried the deletion (Fig 3C). These findings demonstrate a strong but imperfect association between the 1-bp deletion and morph type.

We hypothesized that the nine animals described as *yellowbelly* homozygotes but heterozygous for the 1-bp deletion might carry an alternate loss-of-function variant in *EDNRB1* on the homologous chromosome. To test this hypothesis, we sequenced the coding regions and adjacent splice sites of *EDNRB1* in one of these animals. We observed that this animal was heterozygous for a 17-bp deletion in the fourth coding region of *EDNRB1* (OP589186:c.1747_1763del, Fig 3B). This deletion was located on the homologous chromosome compared to the 1-bp deletion (S1 Fig). The 17-bp deletion introduces a frameshift in the transcript, which removes or alters two of the seven transmembrane helices of the EDNRB1 protein (Fig 3F). The 17-bp deletion thus represents a second putative loss-of-function allele of *EDNRB1*.

To assess the contribution of the 17-bp deletion to phenotypes attributed to the *yellowbelly* allele, we genotyped all animals in our panel for the 17-bp deletion. We found that the 17-bp deletion was present exclusively in the nine *yellowbelly* homozygotes heterozygous for the 1-bp deletion (Fig 3C). In each case, the 17-bp deletion was carried on the homologous chromosome compared to the 1-bp deletion (S2 Table). These results show that every animal described as a *yellowbelly* homozygote was either homozygous for the 1-bp deletion or compound heterozygous for the 1-bp deletion and the 17-bp deletion (Fig 3D). The absence of animals homozygous for the 17-bp deletion may simply reflect the low frequency of this allele in our sample. These data suggest that the allele known to breeders as *yellowbelly* represents the collection of two molecular alleles of *EDNRB1*: a 1-bp deletion and a 17-bp deletion. We propose that these deletions produce the same phenotype and were therefore not recognized by breeders as distinct.

### The *spark* and *specter* alleles are associated with *EDNRB1* missense variants

Animals described as carrying the *spark* or *specter* alleles of the Yellowbelly series show a more moderate loss of coloration than animals described as *yellowbelly* homozygotes (Fig 1B). We therefore predicted that the *spark* and *specter* alleles might represent variants in *EDNRB1* that reduce gene function to a lesser degree than do the 1-bp and 17-bp deletions. To test this hypothesis, we sequenced the coding regions and adjacent splice sites of *EDNRB1* in two animals: one animal described as a *spark/yellowbelly* compound heterozygote and one animal described as a *specter* homozygote. (Our sample did not include any animals described as *spark* homozygotes.) We observed that the *spark/yellowbelly* animal was heterozygous for a missense variant in the first coding region of *EDNRB1* (OP589186:c.G481C). This variant leads to a leucine-to-phenylalanine exchange at position 152 of the protein, termed hereafter L152F (Fig 4C). This animal was also heterozygous for the 1-bp deletion associated with the *yellowbelly* allele, as expected. We observed that the *specter* homozygote was homozygous for a missense variant in the fifth coding region of *EDNRB1* (OP589186:c.G2601C). This variant leads to an arginine-to-proline exchange at position 315 of the protein, termed hereafter R315P (Fig 4C). No other coding or splice-site variants were observed in either animal. Both missense variants affected conserved sites of the EDNRB1 protein, suggesting that they likely disrupt protein function. Position 152, which is located in a transmembrane helix (Fig 4E), is conserved as a leucine or isoleucine across species (Fig 4G). Position 315, which is located in an intracellular loop (Fig 4F), is conserved as an arginine across species (Fig 4G). We conclude that *L152F* and *R315P* are candidates for the *spark* and *specter* alleles, respectively.

**Fig 4.**
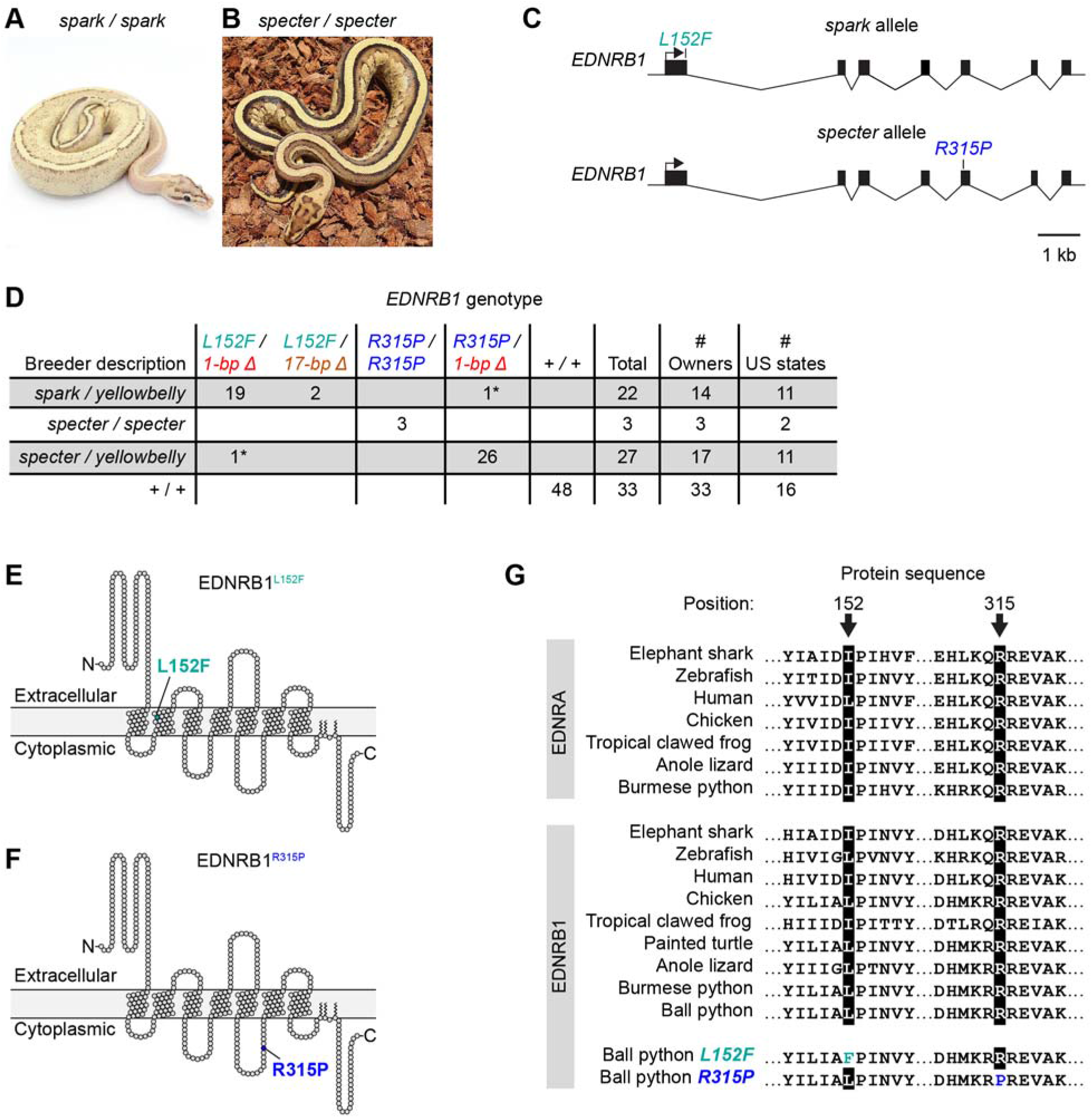
The *spark* and *specter* alleles are associated with missense variants in *EDNRB1*. (A-B) Ball pythons described as homozygous for the *spark* allele or the *specter* allele. Photos, Freedom Breeder, Patrick Faulkner. (C) Schematic of molecular *EDNRB1* alleles associated with the *spark* and *specter* alleles. The *spark* allele contains missense variant *L152F*. The *specter* allele contains missense variant *R315P*. (D) Table showing associations between the *spark* and *specter* alleles and the missense variants. This table includes the original *spark/yellowbelly* compound heterozygote and the original *specter* homozygote in which the missense variants were identified. Numbers of owners and US states indicate the total numbers of owners and US states from which the animals were collected. *, presumed mis-labeling by owner (see Discussion). (E-F) Schematics of EDNRB1 proteins containing missense variants, created using Protter (Omasits *et al*. 2014). (G) Alignment of EDNRB1 and EDNRA protein sequences surrounding missense variants. EDNRA is included to demonstrate conservation outside EDNRB1. Full EDNRA sequences were not available for ball python or painted turtle.

To test whether *L152F* and *R315P* were associated with phenotypes attributed to the *spark* and *specter* alleles, we genotyped these variants in the following panel of animals: 21 additional animals described as *spark/yellowbelly* compound heterozygotes; two additional animals described as *specter* homozygotes; 27 animals described as *specter/yellowbelly* compound heterozygotes; and 48 Non-Yellowbelly animals. Animals were also genotyped for the 1-bp and 17-bp deletions associated with the *yellowbelly* allele. We observed that the missense variants were nearly perfectly associated with the *spark* and *specter* alleles: (i) both animals described as *specter* homozygotes were homozygous for *R315P*, (ii) all but one animal described as *spark/yellowbelly* were heterozygous for *L152F* and for one of the deletions, and (iii) all but one animal described as *specter/yellowbelly* were heterozygous for *R315P* and for one of the deletions (Fig 4D). The exceptions were that one animal described as *spark/yellowbelly* carried *R315P*, rather than *L152F*, and one animal described as *specter/yellowbelly* carried *L152F*, rather than *R315P* (Fig 4D). These exceptions may represent mis-labeling by owners, given the phenotypic similarity between the *spark* and *specter* alleles (see Discussion). These data largely support the hypothesis that *L152F* represents the *spark* allele and *R315P* represents the *specter* allele.

### The *gravel* and *asphalt* alleles are associated with variants in *EDNRB1* splice donors

Animals described as carrying the *gravel* or *asphalt* alleles of the Yellowbelly series show the mildest loss of coloration of all morphs in the series (Fig 1B). We therefore predicted that the *gravel* and *asphalt* alleles might represent variants in *EDNRB1* with mild effects on gene function. To test this hypothesis, we sequenced the coding regions and adjacent splice sites of *EDNRB1* in one animal described as a *gravel* homozygote and one animal described as an *asphalt* homozygote. We observed that both animals were homozygous for synonymous G-to-A nucleotide substitutions in the -1 positions of splice donors. For the *gravel* homozygote, the substitution was located in the splice donor for the first intron (OP589186:c.G499A, Fig 5C). For the *asphalt* homozygote, the substitution was located in the splice donor for the sixth intron (OP589186:c.G3118A, Fig 5C). No coding variants and no other splice-site variants were observed in either animal. The -1 position of splice donors contributes to splicing efficiency across eukaryotes (Cartegni *et al*. 2002), and this position is a guanine in ∼80% of splice donors in vertebrate genes (Ma *et al*. 2015). In *EDNRB1*, the -1 positions of the first and sixth splice donors are conserved as guanines across species (Fig 5C), suggesting that guanines at these sites contribute to gene function. We conclude that the splice-donor substitutions identified herein are candidates for the *gravel* and *asphalt* alleles.

**Fig 5.**
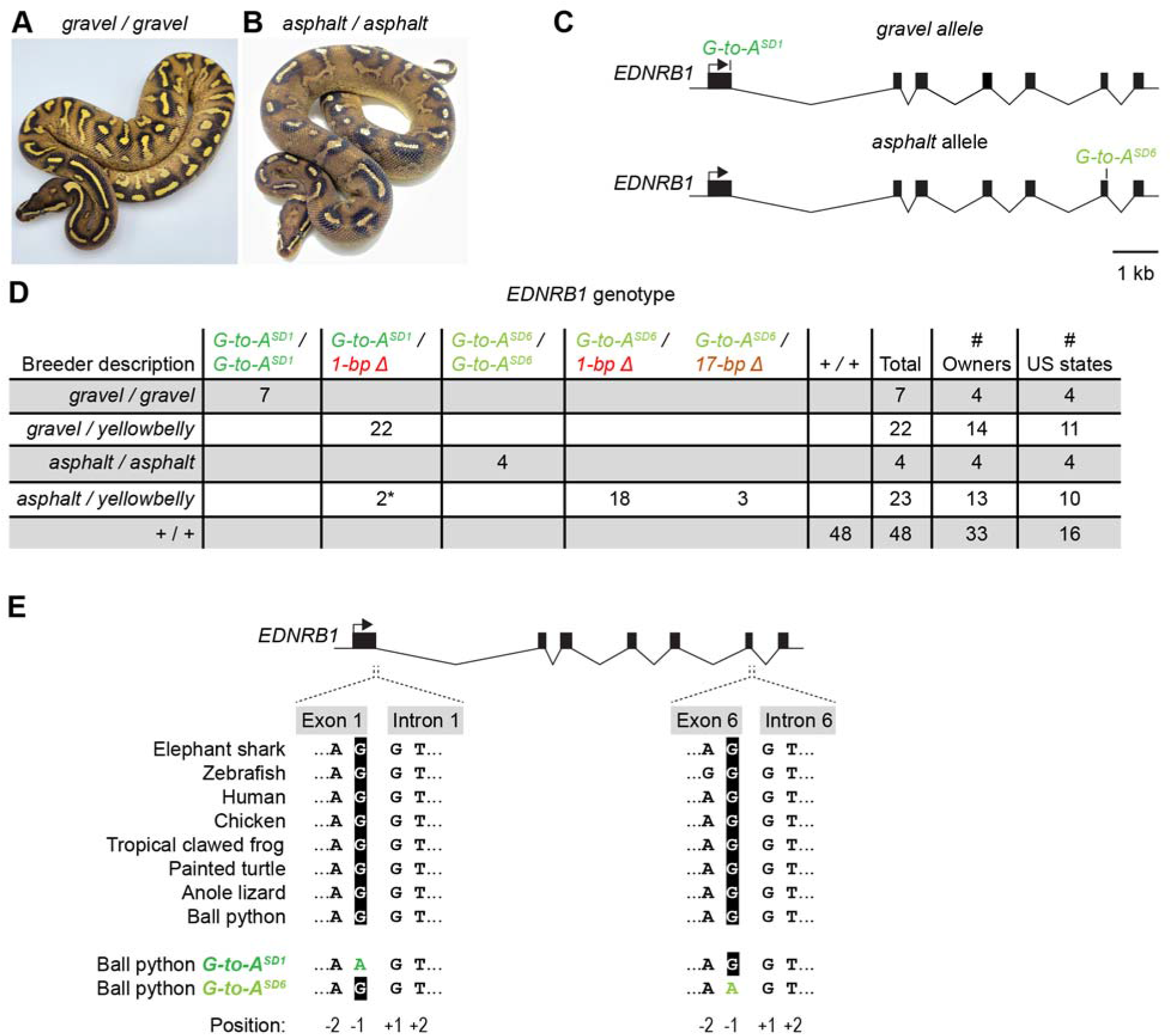
The *gravel* and *asphalt* alleles are associated with nucleotide substitutions in *EDNRB1* splice donors. (A-B) Ball pythons described as homozygous for the *gravel* allele or the *asphalt* allele. Photos, Off the Wall Balls, Ball Python Shed. (C) Schematic of molecular *EDNRB1* alleles associated with the *gravel* and *asphalt* alleles. The *gravel* and *asphalt* alleles contain G-to-A nucleotide substitution in the -1 positions of the first and sixth splice donors, respectively (SD1 and SD6). (D) Table showing associations between the *gravel* and *asphalt* alleles and the nucleotide substitutions in splice donors. This table includes the original *gravel* and *asphalt* homozygotes in which the nucleotide substitutions were identified. Numbers of owners and US states indicate the total numbers of owners and US states from which the animals were collected. *, presumed mis-labeling by owner (see Discussion). (E) Alignment of nucleotide sequences for the first and sixth splice donors of *EDNRB1*.

To test whether the G-to-A substitutions in splice donors were associated with phenotypes attributed to the *gravel* and *asphalt* alleles, we genotyped these variants in the following panel of animals: six additional animals described as *gravel* homozygotes; three additional animals described as *asphalt* homozygotes; 22 animals described as *gravel/yellowbelly* compound heterozygotes; 23 animals described as *asphalt/yellowbelly* compound heterozygotes; and 48 Non-Yellowbelly animals. These animals were also genotyped for the 1-bp and 17-bp deletions associated with the *yellowbelly* allele. We observed that the G-to-A substitutions in splice donors were nearly perfectly associated with the *gravel* and *asphalt* alleles: (i) all animals described as *gravel* or *asphalt* homozygotes were homozygous for the substitutions in the first or sixth splice donors, respectively, (ii) all animals described as *gravel/yellowbelly* were heterozygous for the substitution in the first splice donor and for one of the deletions, and (iii) all but two animals described as *asphalt/yellowbelly* were heterozygous for the substitution in the sixth splice donor and for one of the deletions (Fig 5D). The exceptions were two animals described as *asphalt/yellowbelly*, which the substitution in the first splice donor, rather than in the sixth splice donor (Fig 5D). These exceptions may represent mis-labeling by owners, given the phenotypic similarity between the *gravel* and *asphalt* alleles (see Discussion). These data largely support the hypothesis that the G-to-A substitution in the first splice donor represents the *gravel* allele and the G-to-A substitution in the sixth splice donor represents the *asphalt* allele.

### No candidate variants in *EDN3* or *EDNRB2*

Our initial focus on *EDNRB1*, rather than *EDN3* and *EDNRB2*, relied on sequences from a single *yellowbelly* homozygote. To further exclude *EDN3* and *EDNRB2* as causative for phenotypes in the Yellowbelly series, we amplified and sequenced the coding regions and adjacent splice sites of *EDN3* and *EDNRB2* in the following animals: One *specter* homozygote (*R315P/R315P*), one *gravel* homozygote (*A/A* at the first splice donor), one *asphalt* homozygote (*A/A* at the sixth splice donor), and one *spark/yellowbelly* compound heterozygote (*L152F/1-bp deletion*). We observed no coding variants and no splice-site variants in any of these animals. These results further support the conclusion that phenotypes in the Yellowbelly series are caused by variants in *EDNRB1*, not *EDN3* or *EDNRB2*.

### Reliability of heterozygote descriptions

Ball pythons described as heterozygous for alleles in the Yellowbelly series sell for higher prices than wildtype animals. This price difference provides an incentive for sellers to label animals as heterozygotes. Yet heterozygotes for alleles in the Yellowbelly series are difficult to distinguish from wildtype. Heterozygotes have subtly lighter coloration and altered patterning in the underbelly (Figs 1E and 1F), but this phenotype is variable and overlaps with the range of phenotypes observed among wildtype animals. Thus, animals described as heterozygous for alleles in the Yellowbelly series are viewed with a degree of skepticism. Many believe that some of these animals are mis-labeled, perhaps for monetary gain by sellers.

Our discovery of association between alleles in the Yellowbelly series and variants in *EDNRB1* allowed us to test the reliability of heterozygote descriptions. To do so, we genotyped 130 animals described as *yellowbelly* heterozygotes for the 1-bp and 17-bp deletions in *EDNRB1*. These animals were collected from a total of 68 pet owners across the United States and therefore represent a broad sampling of the pet population. We focused on the *yellowbelly* allele, rather than other alleles in the Yellowbelly series, because the *yellowbelly* allele is described as producing the strongest phenotype in heterozygotes. We observed that 98 of the 130 animals in our sample were heterozygous for the 1-bp or 17-bp deletions in *EDNRB1* (Fig 3C). The animals lacking these deletions were further tested to determine whether they carried any of the other variants associated with alleles in the Yellowbelly series—*L152F* (*spark*), *R315P* (*specter*), and the G-to-A substitutions in splice donors (*gravel* and *asphalt*). One animal was found to be heterozygous for *L152F* (S2 Table). The other animals did not carry any of these variants (S2 Table). Thus, of the 130 animals described as *yellowbelly* heterozygotes, 98 were correctly described, one was a *spark* heterozygote, and the remainder carried only putatively wildtype copies of *EDNRB1*. These findings confirm the skepticism surrounding heterozygote descriptions for alleles in the Yellowbelly series. They suggest that approximately one in four animals described as a *yellowbelly* heterozygote is mis-labeled.

## Discussion

The goal of the current study was to identify the genetic cause of phenotypes in the Yellowbelly series of ball pythons. These phenotypes were characterized by a loss of coloration and changes in patterning, ranging from severe (all-white skin) to moderate (dorsal striping) to mild (subtle changes in patterning compared to wildtype) (Fig 1). Through careful choice of candidate genes and a bit of luck, we discovered that these phenotypes were associated with six putative loss-of-function variants in the gene encoding endothelin receptor EDNRB1 (Fig 6). All-white skin was associated with frameshift variants (Fig 3). Dorsal striping was associated with missense variants (Fig 4). Subtle changes in patterning were associated with splice-donor variants (Fig 5). To our knowledge, these variants represent the first description of genetic variants affecting endothelin signaling in a non-avian reptile.

**Fig 6.**
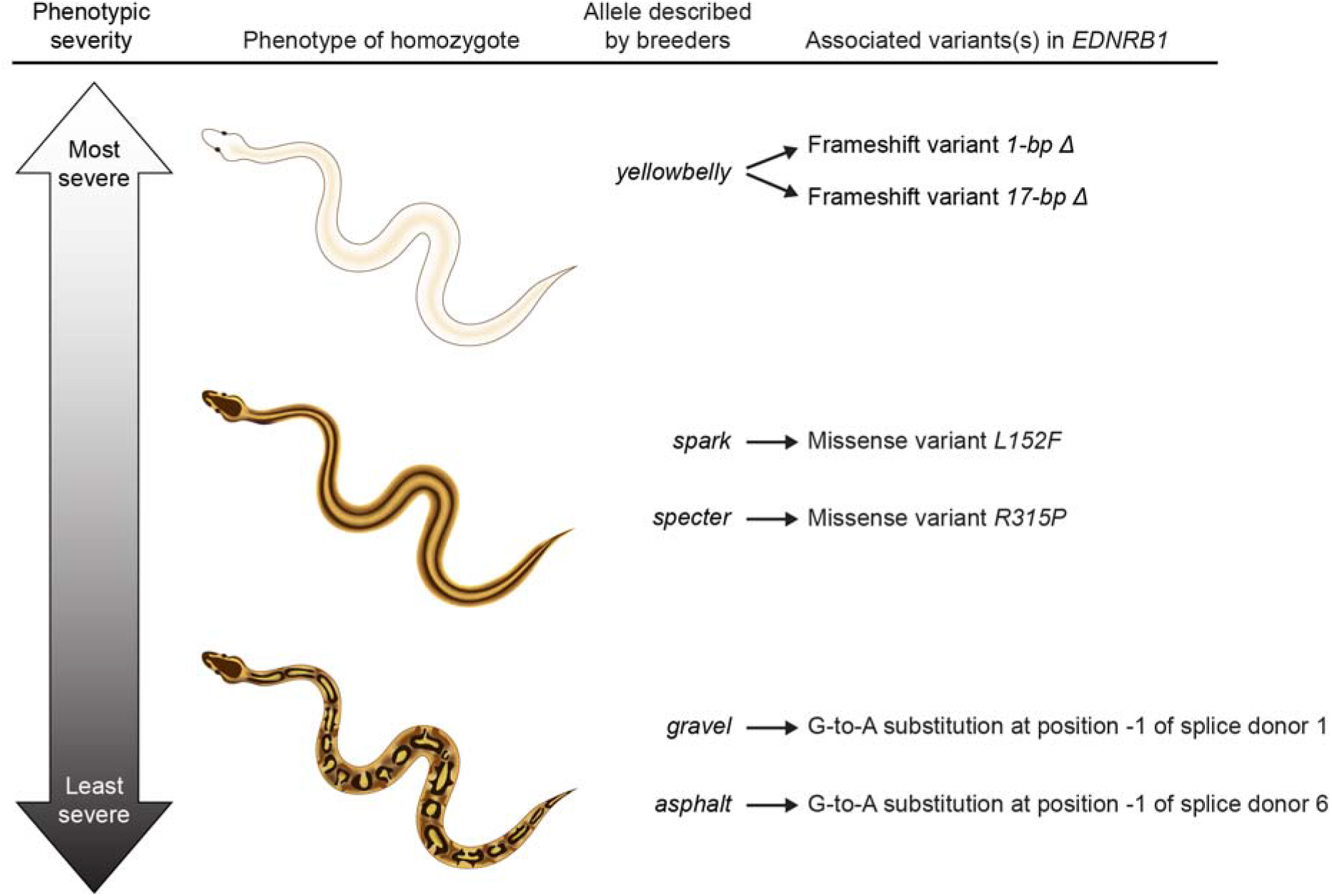
Summary of genetic variants in *EDNRB1* associated with phenotypes in the Yellowbelly series. The most severe phenotype in the Yellowbelly series was all-white skin, which was associated with frameshift variants (1-bp and 17-bp deletions). The next most severe phenotype was dorsal striping, which was associated with missense variants *L152F* and *R315P*. The mildest phenotype was altered patterning, which was associated with G-to-A nucleotide substitutions in the -1 positions of splice donors.

### Causality, limitations, and the candidate-gene approach

We propose that the *EDNRB1* variants described herein are causal for phenotypes in the Yellowbelly series. This idea is supported by a correlation between the severity of phenotypes in the Yellowbelly series and the severity of genetic lesions in *EDNRB1* (Fig 6). The most severe phenotype in the series—all-white skin—was associated with frameshift variants in *EDNRB1*, which plausibly represent null alleles (Fig 3). White skin is characteristic of an absence of melanophores and xanthophores (e.g. Woodcock *et al*. 2017; Ullate-Agote and Tzika 2021), and the association between all-white skin and *EDNRB1* in ball pythons is consistent with *EDNRB1* controlling chromatophore development throughout vertebrates (reviewed in Braasch and Schartl 2014). We propose that all-white skin represents the null phenotype for *EDNRB1* in ball pythons.

The next most severe phenotype in the Yellowbelly series was dorsal striping. This phenotype was associated with missense variants at conserved sites of the EDNRB1 protein (Fig 4). We propose that these variants represent hypo-morphic alleles. These missense variants might reduce EDNRB1 protein function via effects on folding or interactions with co-factors, among other possibilities. Dorsal striping in ball pythons resembles mutant phenotypes in zebrafish caused by iridophores and melanophores being reduced in number and remaining clustered near the horizontal midline (Fig 2A and 2B) (Parichy *et al*. 2000; Lopes *et al*. 2008; Patterson and Parichy 2013; Frohnhöfer *et al*. 2013; Mo *et al*. 2017; Spiewak *et al*. 2018). We propose that stripe formation in ball pythons is analogous to this phenotype in zebrafish, and that stripes in ball pythons are caused by chromatophores remaining clustered near the dorsal ridge (Fig 2E). This phenotype might arise from reduced cell proliferation, reduced cell migration, or both, given that endothelin signaling controls one or both processes in other species (Lahav *et al*. 1996; Reid *et al*. 1996; Pla *et al*. 2005; Harris *et al*. 2008; Krispin *et al*. 2010; Kawasaki-Nishihara *et al*. 2011; Woodcock *et al*. 2017).

The mildest phenotype in the Yellowbelly series was altered patterning. This phenotype was associated with G-to-A nucleotide substitutions in the -1 positions of *EDNRB1* splice donors (Fig 5). The -1 position of splice donors contributes to splicing efficiency (Cartegni *et al*. 2002), and variants in the -1 positions of splice donors of human genes have been known to disrupt gene function (Anna and Monika 2018). We propose that the splice-donor substitutions in *EDNRB1* reduce the fidelity of splicing and thereby reduce EDNRB1 protein levels. The outcome may be a mild reduction in chromatophore number or altered cell migration or both, leading to changes in patterning (Fig 2D). The idea that mild changes cell number or migration could produce noticeable changes in patterning is consistent with the idea that patterns arise via a complex network of cell-to-cell interactions (Singh and Nuesslein-Volhard 2015; Patterson and Parichy 2019), which are sensitive to cell numbers (Spiewak *et al*. 2018) and other parameters (Watanabe and Kondo 2012; Kaelin *et al*. 2012, 2021; McGuirl *et al*. 2020).

The scope of our study was limited to three genes: *EDN3, EDNRB1* and *EDNRB2*. Thus, we cannot formally exclude the possibility that variants in other regions of the genome contribute to phenotypes in the Yellowbelly series. However, we view this possibility as unlikely, given that six independent haplotypes of *EDNRB1* were associated with phenotypes in the Yellowbelly series, and that the severity of these phenotypes correlated with the severity of genetic lesions in *EDNRB1*. We propose that further evidence supporting a link between *EDNRB1* and the Yellowbelly series could be obtained by introducing the variants described herein into a model system (e.g. zebrafish) to test for effects on protein function or gene expression.

An additional limitation of our study is that samples were limited to shed skins. Shed skins did not allow us to compare chromatophore numbers across morphs, nor did they allow us to characterize chromatophore behavior during development. Characterization of embryos from wildtype ball pythons, as well as embryos in the Yellowbelly series, is needed to fully characterize the role of *EDNRB1* in ball pythons.

### Pros and cons of community-sourced sampling

Our study relied on pet samples recruited from the community, rather than samples from a single pedigree. An advantage of this approach was increased genetic diversity in our sample. An example of this diversity was the discovery that the allele known to breeders as *yellowbelly* represents two distinct molecular alleles of *EDNRB1*—a 1-bp deletion and a 17-bp deletion (Fig 3). The 17-bp deletion was approximately 16-fold less common in our sample than the 1-bp deletion and might have been plausibly missed by our study, had we focused on a single pedigree. The phenomenon of multiple molecular alleles corresponding to a single breeder-defined allele is not unique to the Yellowbelly series and has been reported for two other color morphs in ball pythons—the Albino color morph and the Ultramel color morph (Brown *et al*. 2022). This phenomenon highlights the utility of community-sourced sampling in ball pythons and shows that the pet population is more diverse than previously recognized.

A disadvantage of using pet samples sourced from the community is that genotype descriptions provided by owners come with a degree of uncertainty. Certain combinations of alleles in the Yellowbelly series produce similar phenotypes (e.g. *spark* vs. *specter* and *gravel* vs. *asphalt*). Crosses involving these alleles therefore generate progeny whose genotypes are uncertain. Breeders typically label such animals according to their best guess of the animal’s genotype. This practice leads to mis-labeling by owners and likely explains the apparent mix-ups of animals in our study: one *spark/yellowbelly* animal was mis-labeled by its owner as *specter/yellowbelly* and vice versa (Fig 4D); two *gravel/yellowbelly* animals were mis-labeled by their owners as *asphalt/yellowbelly* (Fig 5D); and one *spark* heterozygote was mis-labeled by its owner as a *yellowbelly* heterozygote (Fig 3C). An additional cause of mis-labeling is that breeders are incentivized to label animals as heterozygous for alleles in the Yellowbelly series, even when the phenotype of heterozygotes is difficult to recognize (Figs 1E and 1F). This incentive likely explains why approximately one in four animals described as *yellowbelly* heterozygotes carried only putatively wildtype copies of *EDNRB1* (Fig 3C). We propose that mis-labeling can be reduced in the future by genotyping animals for the *EDNRB1* variants reported in our study.

### Evolution of endothelin signaling and stripe formation

The phenotypes associated with *EDNRB1* variants in ball pythons suggest that endothelin signaling may be required for the development of melanophores, xanthophores, and possibly iridophores in this species. Endothelin signaling in snakes may therefore play a role similar to endothelin signaling in axolotl, where the endothelin ligand EDN3 is thought to direct the development of all three types of chromatophores (Woodcock *et al*. 2017). Similar regulation has been described for mammals and birds, where endothelin signals regulate the development of melanophores, which are the only type of chromatophore present in these taxa (Saldana-Caboverde and Kos 2010; Bondurand *et al*. 2018). These species contrast with zebrafish, where endothelin signaling directly promotes the development of iridophores, but is dispensable for xanthophores and plays only an indirect role for melanophores (Patterson and Parichy 2013; Frohnhöfer *et al*. 2013; Patterson *et al*. 2014). These observations have led to the proposal that endothelin signaling in vertebrates was ancestrally required for all three types of chromatophores and became specialized for iridophores in teleost fish (Spiewak *et al*. 2018). Our findings support this hypothesis by adding ball pythons to the list of species in which endothelin signaling likely regulates the development of multiple types of chromatophores.

A notable difference between ball pythons and mammals is that loss of *EDNRB1* function in mammals is lethal. *EDNRB1* in mammals is required for multiple lineages of the neural crest, including chromatophores and enteric neurons of the hindgut (Saldana-Caboverde and Kos 2010; Bondurand *et al*. 2018). Loss of these neurons prevents passage of food through the intestine, causing death of *EDNRB1* mutants in mammals shortly after birth (Hosoda *et al*. 1994; Puffenberger *et al*. 1994; Gariepy *et al*. 1996; Santschi *et al*. 1998; Yang *et al*. 1998; Lühken *et al*. 2012). Surprisingly, morphs in the Yellowbelly series do not exhibit this defect. Even morphs homozygous for presumed null alleles of *EDNRB1* are viable and healthy, at least in captivity. One possible explanation for this viability is that *EDNRB1* in ball pythons may be specialized for chromatophore lineages of the neural crest. Neuronal lineages might use the other B-type endothelin receptor, *EDNRB2*. This specialization is not possible in mammals because eutherian mammals have lost *EDNRB2*, owning to its location on the chromosomes that gave rise to the mammalian sex chromosomes (Braasch *et al*. 2009). Specialization of B-type endothelin receptors may also have occurred along the lineage leading to birds, but in the opposite direction. White plumage in several bird species has been associated with loss-of-function variants in *EDNRB2*, but never *EDNRB1* (Miwa *et al*. 2007; Kinoshita *et al*. 2014; Li *et al*. 2015; Xi *et al*. 2020). Thus, our identification of *EDNRB1* variants in ball pythons as associated with color phenotypes provides insight into the evolution of endothelin receptors throughout higher vertebrates.

Dorsal, longitudinal stripes are not unique to morphs in the Yellowbelly series. Other striped morphs in ball pythons include the Genetic Stripe morph, the Champagne morph, and some morphs in the Blue-Eyed Leucistic series (McCurley 2005; Broghammer 2019). Outside ball pythons, stripes occur naturally in many snake species, typically in species that escape predation via rapid flight (Allen *et al*. 2013). Stripes have arisen multiple times within the snake lineage (Wolf and Werner 1994), and a few species are naturally polymorphic for striped and non-striped forms (Zweifel 1981; Brodie 1992; King 2003; Westphal and Morgan 2010; Kuriyama *et al*. 2013; Murakami *et al*. 2014). Whether stripe formation in these cases involves changes to endothelin signaling or chromatophore number remains to be determined. Our study suggests that dorsal, longitudinal stripes may be easily accessible to the snake body form via genetic changes that alter simple developmental parameters in the neural crest, such as cell number. Candidate genes for controlling these parameters include other genes in the endothelin pathway (Braasch *et al*. 2009) and genes responsible for specifying cell fates in the neural crest (Parichy and Spiewak 2015; Mort *et al*. 2015).

## Methods

### Recruitment of ball python sheds

Ball python sheds were recruited from pet owners and breeders by placing announcements in Twitter, Reddit, Instagram, and Facebook, and by contacting sellers having active listings on Morph Market (www.morphmarket.com). Contributors were instructed to allow sheds to dry (if wet) and to ship sheds via standard first-class mail. Contributors sending multiple sheds were instructed to package sheds individually in plastic bags during shipping. Contributors were not provided monetary compensation for sheds, although some contributors were given pre-paid shipping envelopes to cover shipping costs. Upon receipt, sheds were stored at -20°C to kill any insect larvae infesting the sheds.

We attempted to maximize genetic diversity within each category of morph by excluding animals described as siblings, parents, or offspring of another animal in our sample. However, we cannot exclude the possibility that some animals in our sample may have been close relatives, given that full pedigree information was not available for most animals. The idea that animals in our sample were derived from multiple lineages is supported by our discovery of multiple molecular variants corresponding to the allele known to breeders as *yellowbelly*. Owner codes and US states of residence for each animal are provided in S2 Table.

The total set of animals used in our study were 70 animals described as *yellowbelly* homozygotes, 130 animals described as *yellowbelly* heterozygotes, 22 animals described as *spark/yellowbelly*, three animals described as *specter* homozygotes, 27 animals described as *specter/yellowbelly*, seven animals described as *gravel* homozygotes, 22 animals described as *gravel/yellowbelly*, four animals described as *asphalt* homozygotes, 23 animals described as *asphalt/yellowbelly*, and 48 animals described as having wildtype coloration or belonging to morphs other than those in the Yellowbelly series (Non-Yellowbelly animals).

### Performing experiments in an undergraduate laboratory course

The majority of experiments and analyses described in this study were performed by undergraduate students as part of a laboratory course at Eastern Michigan University (BIO306W). This practice required that our experimental design rely on simple techniques, namely PCR, restriction digests, and Sanger sequencing. To avoid student errors in these techniques, we implemented the following precautions. First, students never handled more than one shed skin at the same time. Second, students performed negative and positive control reactions for all PCR amplifications and restriction digests. Data from students having incorrect controls were excluded from analysis. Third, all sequence analyses were performed independently by three or more students. When the results of all students did not all agree, sequences were re-analyzed by the instructor (HSS). Fourth, when we encountered an animal whose molecular genotype did not match the breeder description (e.g. the single animal labeled *spark/yellowbelly* whose genotype matched *specter/yellowbelly* and vice versa—see Results and Discussion), DNA from the animal was re-extracted by the instructor (HSS), and the molecular genotype were determined a second time. This precaution was taken to guard against mix-ups by students, but it proved unnecessary because the second round of genotypes always matched the first.

### DNA extraction

Sheds containing visible dirt or debris were rinsed in tap water until clean. Sheds were air dried and lysed overnight at ∼60°C in ∼1 ml lysis buffer (100 mM Tris-HCl pH 8.0, 100 mM EDTA, 2% sodium dodecyl sulfate, 3 mM CaCl2, 2 mg/ml Proteinase K) per palm-sized piece of shed. Lysis reactions were performed in standard 1.5 ml tubes. In some cases, pieces of shed were snipped or torn into multiple pieces to enable them to fit into tubes more easily. Dorsal and ventral scales were used interchangeably, after preliminary studies indicated that DNA could be reliably extracted from both types of scales.

Lysate was separated from visible fragments of undigested shed and further cleared by centrifugation at 13,000 x g for 2 min. To precipitate protein, ammonium acetate was added to supernatant to a final concentration of 1.875 – 2.5 M. Samples were incubated on ice for 5 min and centrifuged at 13,000 x g for 3 min at 4°C. Supernatant was mixed with an equal volume of magnetic bead mixture (10 mM Tris-HCl pH 8.0, 1 mM EDTA, 1.6 M NaCl, 0.2% Tween-20, 11% polyethylene glycol, 0.04% washed SpeedBeads [Sigma #GE45152105050250]) and shaken for 5 min. Beads were separated from supernatant using a magnet, washed twice in 0.2 ml 70% ethanol for 2 min, and air dried for ∼1 min. DNA was eluted from beads in TE buffer (10 mM Tris-HCl pH 8.0, 1 mM EDTA) at 65°C for >5 min.

### Primer design and PCR

Primers were designed against the genome of Burmese python, the closest relative of ball python for which genome sequence was available (Castoe *et al*. 2013). Annotations for Burmese python *EDNRB1* were XM_007436628 (mRNA) and XP_007436690 (protein). Annotations for Burmese python *EDN3* were XM_025169570 (mRNA) and XP_007432959 (protein). These annotations were manually examined for errors, by compared gene structure across vertebrates. For both genes, annotations were determined to be correct, based on the observation that gene structure matched gene structure in other vertebrates, including in species with higher quality annotations, such as mouse, chicken, and zebrafish.

Primers were designed using Primer3 (Untergasser *et al*. 2012), using default parameters and a target annealing temperature of 60°C. Amplification was performed using an annealing temperature of 57°C, to allow for occasional divergence between ball python and Burmese python genomic sequences. Coding regions of *EDNRB1* and *EDN3* were amplified and sequenced using primers given in S1 Table.

PCR was performed using OneTaq polymerase (NEB #M0480). Reactions consisted of 1X OneTaq Standard Reaction Buffer, 200 µM dNTPs, 0.2 µM of each primer, and 0.025 U/µl OneTaq polymerase. Thermocycling conditions were as follows: 94°C for 2 min; 30-35 cycles of 94°C for 30 sec, 57°C for 30 sec, and 68°C for 1-2 min; and 68°C for 5 min. Reactions used 10-100 ng template DNA per 20 µl volume.

### Sanger sequencing

PCR products were purified for Sanger sequencing using magnetic beads. PCR reactions were mixed with three volumes of magnetic-bead mixture (10 mM Tris-HCl pH 8.0, 1 mM EDTA, 1.6-2.5 M NaCl, 0.2% Tween-20, 11-20% polyethylene glycol, 0.04% washed SpeedBeads [Sigma #GE45152105050250]) and agitated for 5 min. Beads were separated from supernatant using a magnet, washed twice in 0.2 ml 80% ethanol for >30 sec, and air-dried for 30 sec. PCR products were eluted from beads in 10 mM Tris-HCl pH 8.0 for >3 min at 65°C. Sanger sequencing was performed by Eton Bioscience Inc (etonbio.com). Sequencing primers are provided in S1 Table.

### Protein sequence alignment

Protein sequences were aligned using Clustal Omega (Sievers *et al*. 2011), using default parameters. Accession numbers for protein sequences were as follows: elephant shark EDNRB1 (XP_007900764) and EDNRA (XP_007897347); zebrafish EDNRBb (XP_688565) and EDNRA (NP_001092915); human EDNRB1 (NP_000106) and EDNRA (NP_001948); chicken EDNRB1 (XP_015133631) and EDNRA (NP_989450); tropical clawed frog EDNRB1 (XP_012812722) and EDNRA (NP_001072275); painted turtle EDNRB1 (XP_005311368); anole lizard EDNRB1 (XP_003223432) and EDNRA (XP_003221737); Burmese python EDNRB1 (XP_007436690) and EDNRA (XP_007426479).

### Genotyping assays

The 1-bp and 17-bp deletions in *EDNRB1* (*yellowbelly*) were genotyped by amplifying the fourth coding region of *EDNRB1* via PCR and digesting the PCR products with the restriction enzyme *Bsr*I (New England BioLabs, #R0527S). Both deletions eliminate cut sites for *Bsr*I and can thus be genotyped simultaneously (S1 Fig). Primers for amplification were p257F (5’-GGT TCC CAG TCT CTG CAC AT-3’) and p257R (5’-TTC AAC CAA CGC ACA TCA TT-3’). Digestions consisted of 1X NEBuffer™ r3.1 and 0.05 U/µl *Bsr*I. Digestions were performed at 65°C for at least 2 hrs.

Missense variant *R315P* (*specter*) was genotyped by amplifying the fifth coding region of *EDNRB1* via PCR and digesting the PCR products with the restriction enzyme *Sma*I (New England BioLabs, #R0141S). The *R315P* variant creates a cut site for *Sma*I (S2 Fig). Primers for amplification were p258F (5’-ATG GAG TGG ACA GGA TGG AA-3’) and p258R (5’-CCA GCT AGG TGG GGT AGA CA-3’). Digestions consisted of 1X rCutSmart™ Buffer and 0.05-0.1 U/µl *Sma*I. Digestions were performed at 25°C for at least 2 hrs.

Missense variant *L152F* (*spark*) and the G-to-A substitution at the first splice donor (*gravel*) were genotyped by amplifying a fragment containing the first coding region of *EDNRB1* using primers p381F (5’-CAC CAT AAT GCT TAA CAC ACA CAA-3’) and p381R (5’-TCA TAG CAA TGT GAT AAA CCC ACT-3’). PCR products were sequenced with p381F (S3 Fig). The G-to-A substitution at the sixth splice donor (*asphalt*) was genotyped by amplifying a fragment containing the sixth and seventh coding regions of *EDNRB1* using primers p259F (5’-GGC AGG AAA ACT GCT CGA TA-3’) and p260R (5’-CAA ACC AAA GTC CCA GCA TC-3’). PCR products were sequenced with p259F (S3 Fig).

## Acknowledgements

We thank Bob Winning, Katy Greenwald, and David Kass for comments on the manuscript and the Educational Course Support program of New England BioLabs for reagents used in undergraduate teaching labs.

Most of the data in this study were collected by undergraduates enrolled in a laboratory course at Eastern Michigan University. These students constitute the BIO306W Consortium. These students were Bayan Abdeljalil, Garrett Bailey, John (Teddy) Belman, Keith Camac, Aaron Ellis, Amber Fatima, Delaney Garcia, Shannon Gregory, Amber Haley, Chloe Harrison, Carly Kosanovich, Corey Melcher, Lindsey Miracle, Natalia Pineda, Catherine Redding, Audrey Salsido, Caitlin Satler, Jake Sealy, Ananya Shukla, Hannah Strasser, Nehul Tanna, Evan Veenhuis, Syed Wasiuddin, Anna Watson, Madyson Weaver, Raymond Wells, and Kathleen Weymouth.

Shed skins were contributed by the following individuals: Abby Kruse, Adam and Nicole Schmid, Adam of Proper Royals, Allen Gage, Alycia Butler, Amanda Hall, Amanda Yereance, Ammar Hanif, Andelyn Czajka, Andrew Lyons, Andy Rose, Anthony of Anthill Python, AR Walters, Bankrupt Reptiles, Barbara Penyak, Big D, Brent McKelvey, Bryan Rivera, Charles Wilson, Chelsey Wheeler, Chiron Graves, Chris Hernan, Chris Shelly, Chun Ku, CPJ Reptiles, Cynthia Jones, Dale Porcher, Dave Dunn of Ball in Hand Pythons, David Burstein, Denise Jones, E & L’s Livin Art Morphs 2020, Epic Vibrant Balls, Eric Chung, Everything’s Better Orange, FNA Balls, Francis X. DiNino Jr, Gavin Costello, Greg and Kathee of Royally Morphadellic, Harry Roman, Jacob Rodenhausen, Jaden Christensen, Jake and Ashley Brewster, Jake Jewis, James and Angie Rehus of JAR Pythons, Jamie Palazzo, Jason Roberts, Jeff Kearns, Jeff Linton, Jeromy Shaffer, Joe and Wendy Ducos, Joe Myers, John White, Jon Bidwell, Jon Braga, Jon Godek, Justin Orbach of Just Incrediballs, Kai Li, Kandie Tucker, Kassandra Royer, Kelsi Greene, Kenneth Evans, Korie Dy, KT Creations, Laura Carter, Lauren Glynn, Leah Alexander, Leviathan Snakes, Lindsey VanOrman, Loren Morales, Lynnet Melton, Margaret Lentz, Maryann Barbon, Matt Huck, Michael Cole, Michael Nino, Michael Osmun, Miranda Joy Turpin, Morgan Evans, Nate Fitzpatrick of Baltimore Balls, Nathan Granoff, Nick Deinhart, Nicole Gallagher, OkieMom Balls, Ozzy Maldonado, Pedro Alequin of Imperial Morphs, Peter Holden, Phil Phillips, Phil Snyder, Randy Remington, Redstrom Reptiles, Reggie’s Urban Jungle, Roger Gray, Royal Black Balls, Ryan Boyd and Brittney Delacruz, Sandra Frye, Sean Strait of The Morph Lab, Sergio McDole, Shomai Sanford, Steven Nordahl, Tammy Hutchinson of Hutches Clutches, The Wagner’s of Texas, Tia Vogel, Tom Harbin, Troy Stallings, Tyler Grant, Wes Mosher, Wilbanks Reptiles, Zac Parpart, and several anonymous contributors.

## Author Contributions

Conceived and designed the experiments: HSS. Performed the experiments: UMD IL RLT BS TBC HSS. Analyzed the data: UMD IL RLT BS TBC HSS. Contributed reagents or materials: CWG HSS. Wrote the paper: HSS.

## Supporting Tables

**S1 Table. Primers used to amplify and sequence coding regions of *EDN3, EDNRB1*, and *EDNRB2***.

**S2 Table. Genotypes of all animals**.

**S1 Fig.**
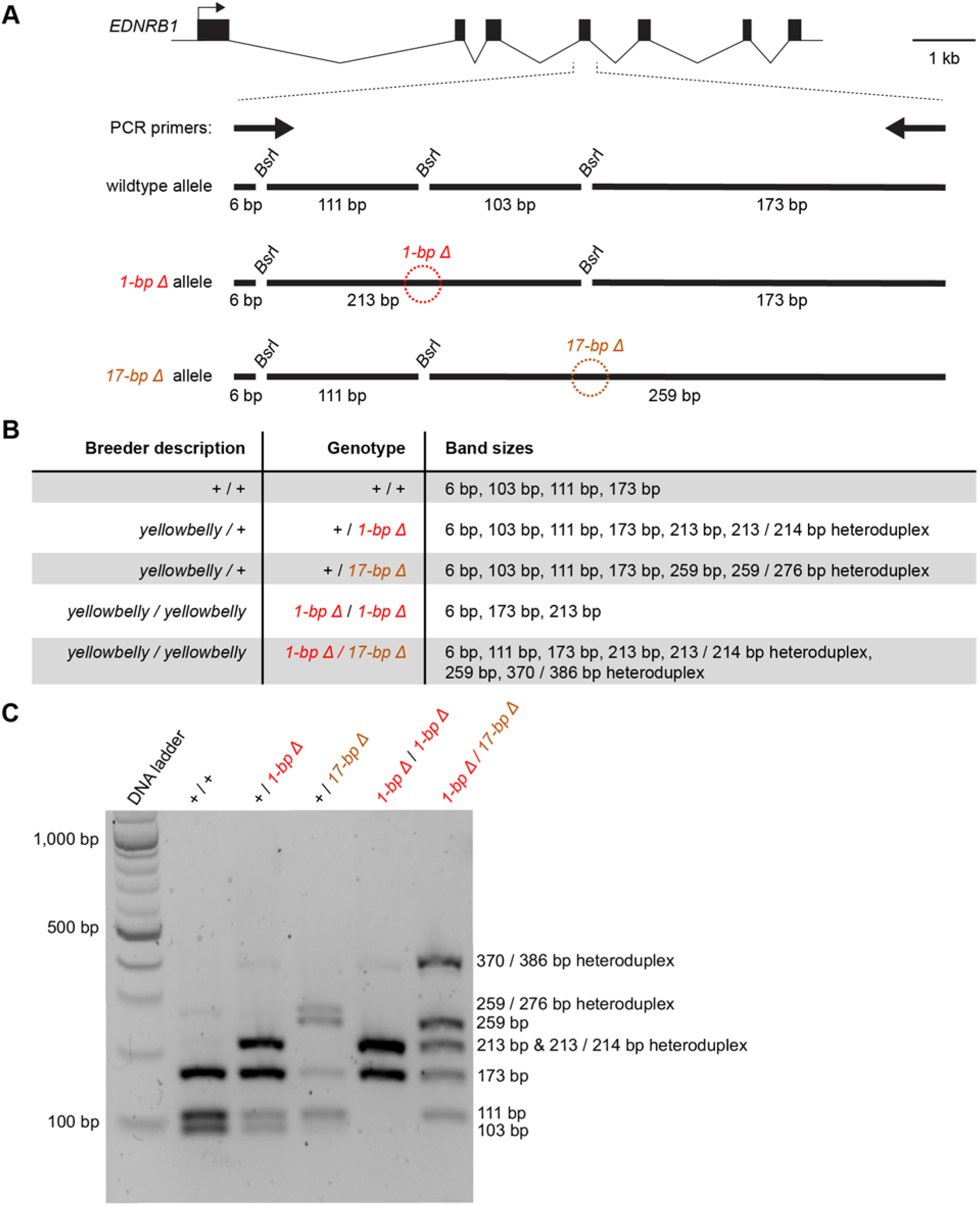
Assay for genotyping the 1-bp and 17-bp deletions in *EDNRB1*. (A) Schematic of the wildtype allele, the 1-bp deletion allele, and the 17-bp deletion allele. The wildtype allele contains three cut sites for *Bsr*I. The deletions each eliminate one of these sites. (B) Band sizes observed for each genotype, demonstrating that the 1-bp and 17-dp deletions were carried on homologous chromosomes. Heteroduplexes were formed because digests were performed at 65**°**C. (C) Agarose gel electrophoresis of *Bsr*I digests.

**S2 Fig.**
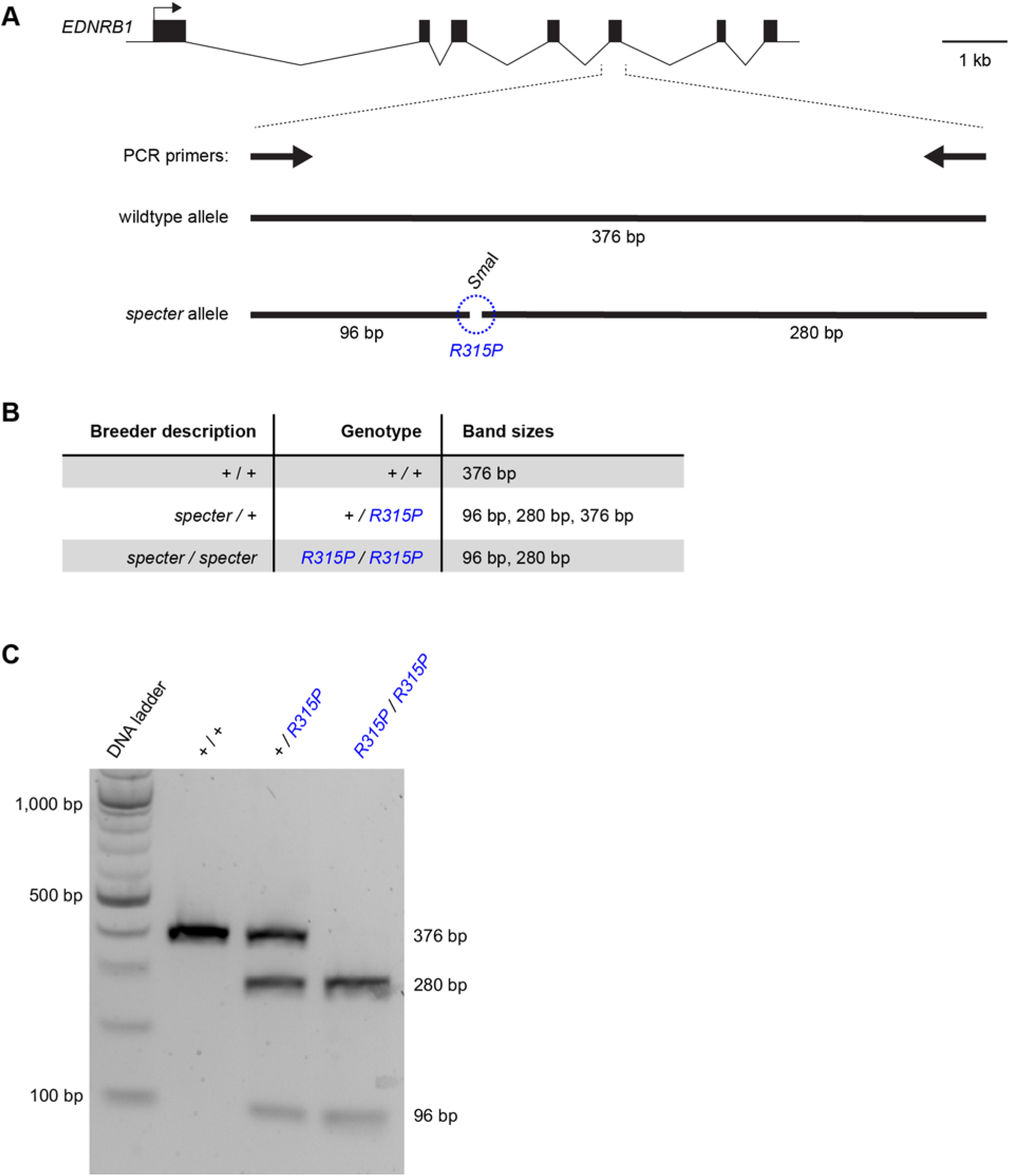
Assay for genotyping the *R315P* missense variant in *EDNRB1*. (A) Schematic of the wildtype allele and the *R315P* allele. The *R315P* variant creates a cut site for *Sma*I. (B) Band sizes observed for each genotype. (C) Agarose gel electrophoresis of *Sma*I digests.

**S3 Fig.**
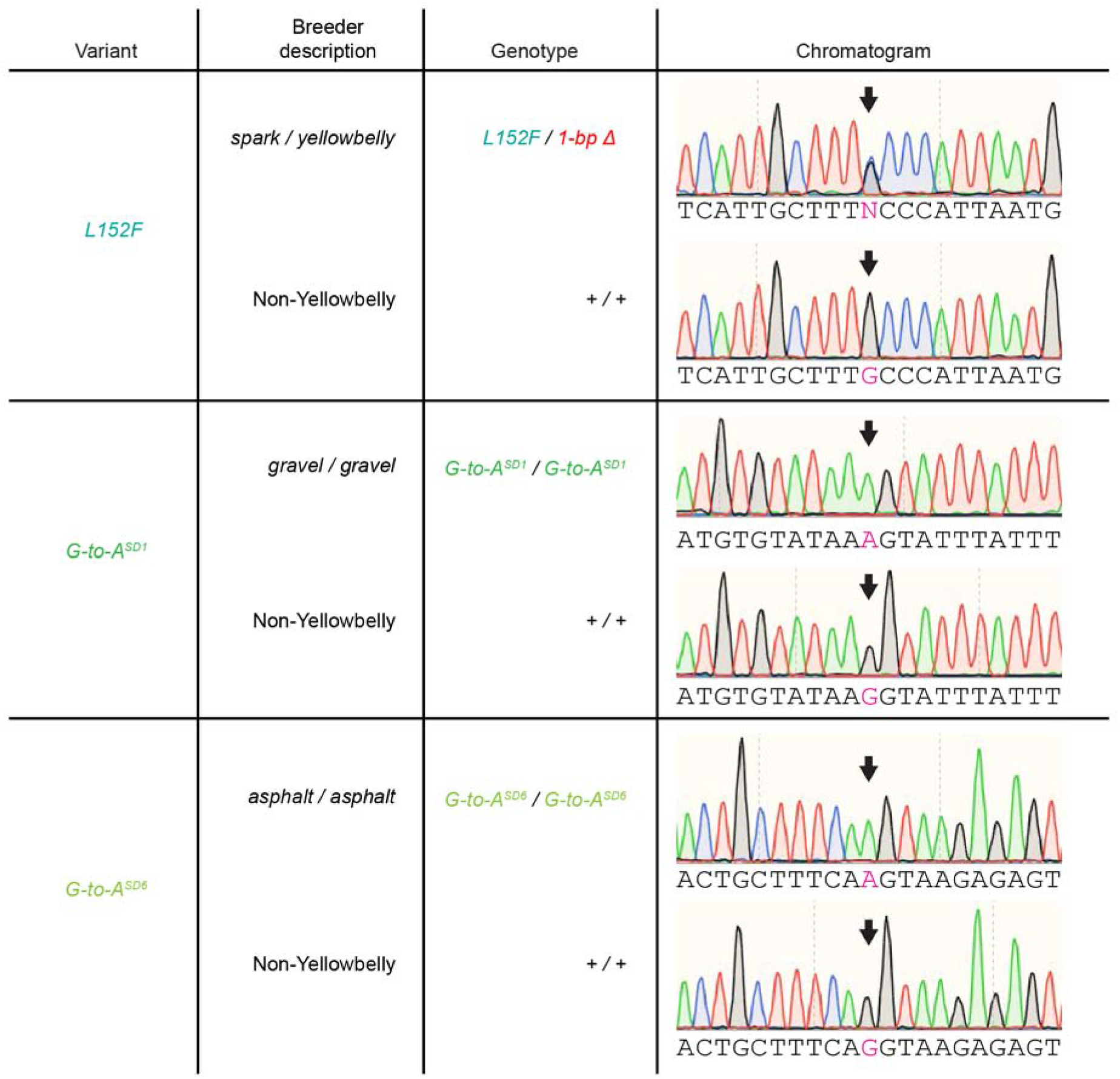
Chromatogram examples. Examples of chromatograms from Sanger sequencing reads used to genotype the *L152F* missense variant and the G-to-A substitutions in splice donors. SD1, splice donor for the first intron. SD6, splice donor for the sixth intron.

