## Supplementary figures and images for "Stripes and loss of color in ball pythons (*Python regius*) are associated with variants affecting endothelin signaling"

### S1_Figure

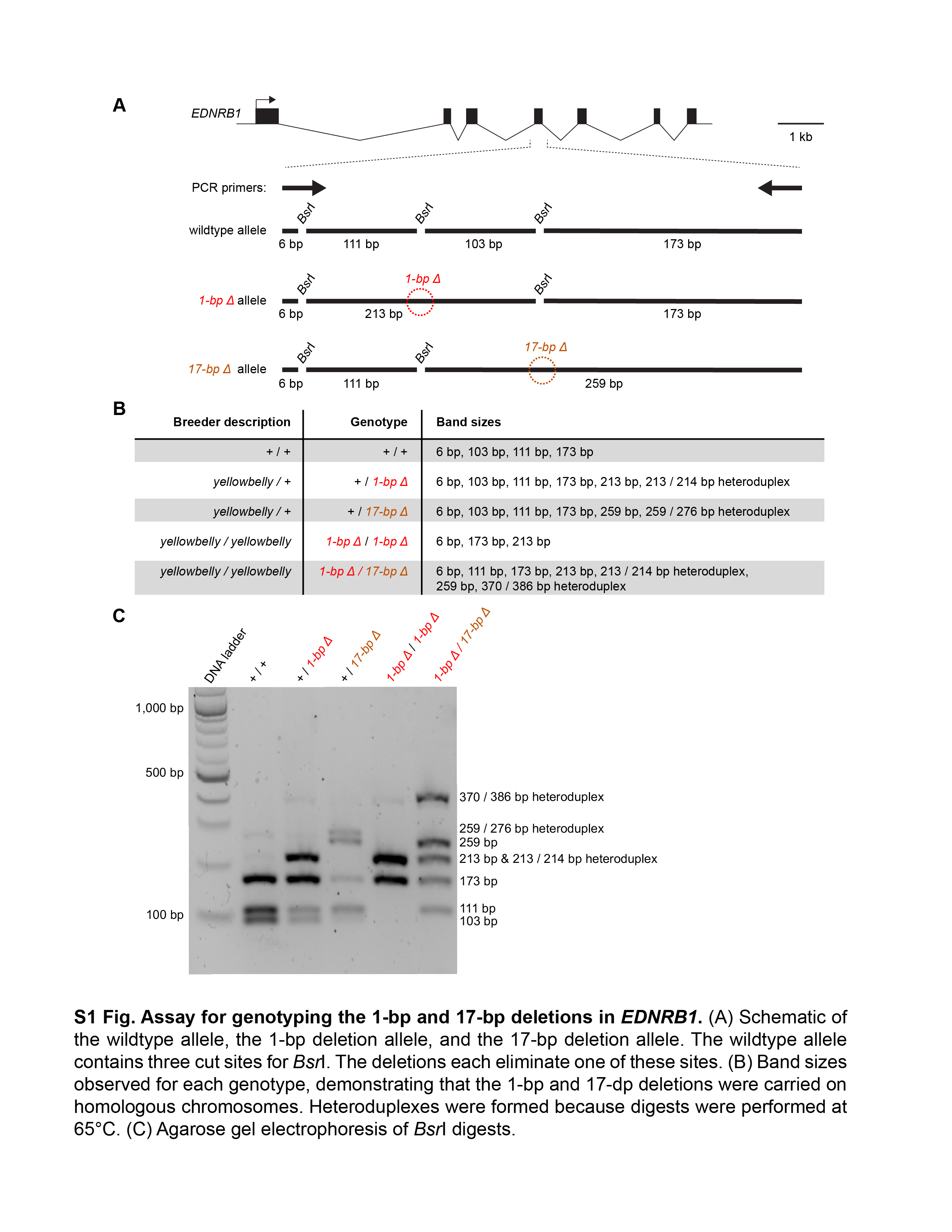

### S2_Figure

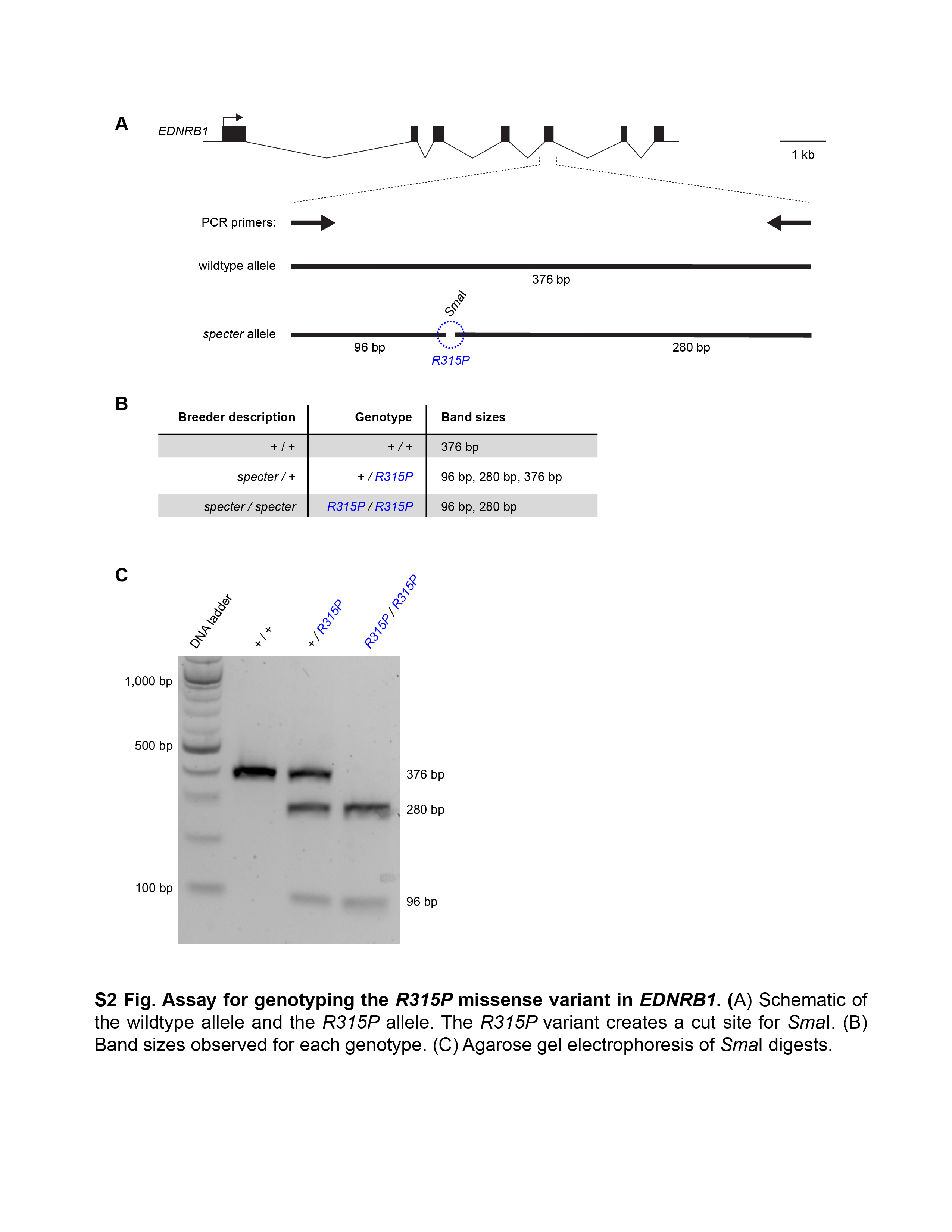

### S3_Figure

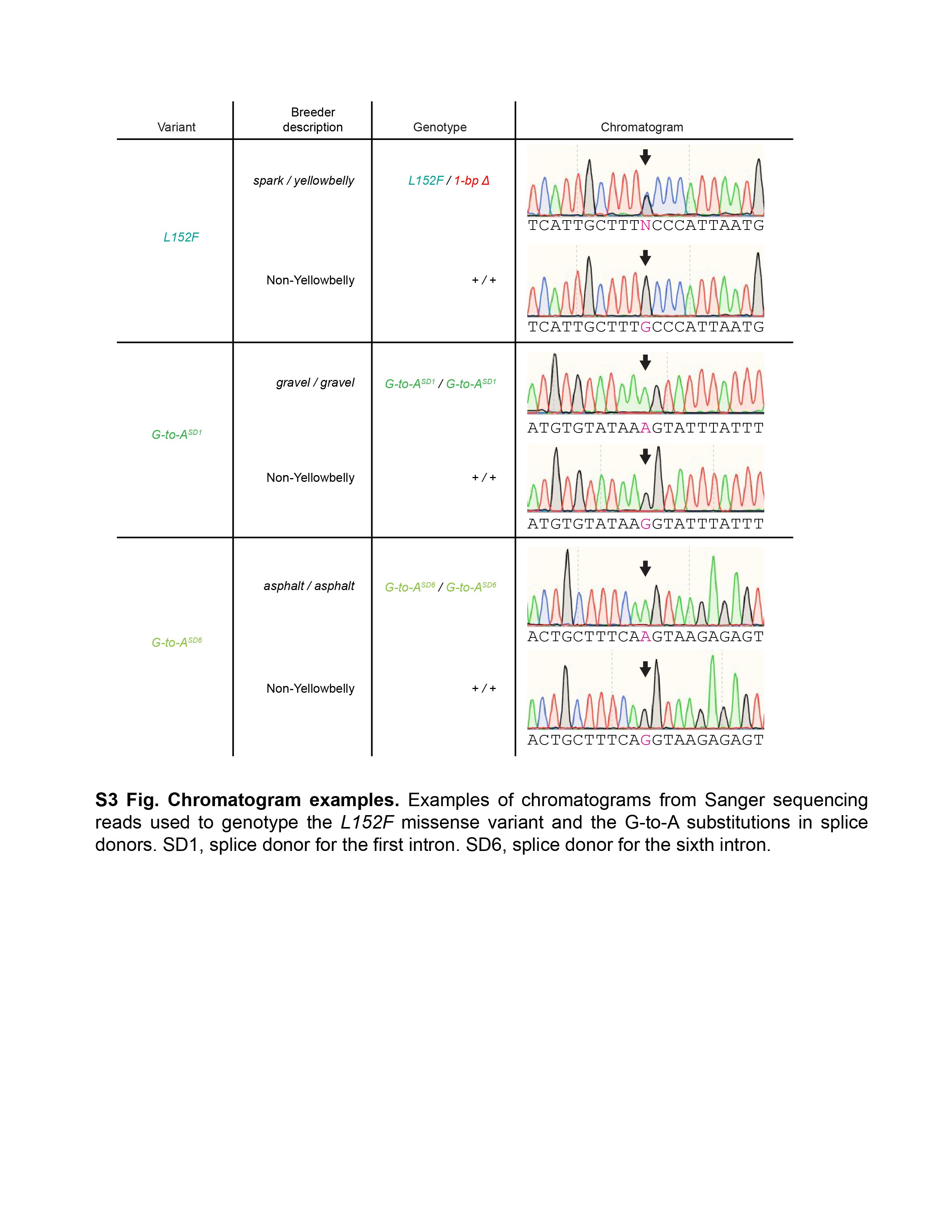
